# Impact of Isolation and Storage Methods on the Properties of Neural Extracellular Vesicles

**DOI:** 10.64898/2026.06.12.731981

**Authors:** Morgane E. Golan, Leah C. McCarthy, Kanupriya R. Daga, Frank Seipel, Randolph S. Ashton, Ross A. Marklein, Steven L. Stice

## Abstract

Extracellular vesicles (EVs) are nanoscale, cell-secreted mediators of intercellular communication with growing promise as therapeutic agents. Manufacturing practices, including EV isolation and storage approaches, are critical determinants of product consistency, purity, and potency. In this study, neural stem cell (NSC)-derived EVs were isolated from conditioned NSC culture media via oscillator-based isolation (OSC), ultracentrifugation (1 or 2 hours), and ultrafiltration, and were stored lyophilized or cryopreserved. Nanoparticle yield, size distribution, and subpopulation composition were evaluated by nano-flow cytometry, quantifying total nanoparticles, membrane-bound EVs and CD63+ EVs. Purification was calculated via particle-to-protein ratios, morphology was evaluated by transmission electron microscopy, and potency was assessed using a microglia morphology assay. Particle yield was comparable across isolation methods, though protein clearance varied, with OSC demonstrating purification relative to conditioned media. Lyophilized samples retained structural integrity, size, and population profiles comparable to cryopreserved samples. Lyophilized and cryopreserved EVs exhibited dose-dependent immunomodulatory activity in our microglia morphology assay, with significant effects observed at 200,000 EVs per cell. These findings highlight the importance of isolation method in EV product quality and support lyophilization as a viable storage strategy which overcomes the logistical limitations of cryopreservation, thereby advancing the development of a robust pipeline for therapeutic EV manufacture.

## INTRODUCTION

Neural stem cell (NSC)-derived extracellular vesicles (EVs) are lipid-bound, cell-secreted nanoparticles with therapeutic potential for treating neurodegenerative diseases and neurotraumatic conditions [1–3]. The clinical translation of EVs hinges upon their ability to deliver bioactive cargoes including proteins, lipids, and RNAs to recipient cells, thereby modulating their behavior and pathological processes [4, 5]. For example, NSC-EVs exhibit context-dependent communication with microglia, demonstrating therapeutic activity in pro-inflammatory neurological states [6]. In fact, in 2024 a US Food and Drug Administration (FDA) Investigational New Drug (IND) application for neural stem cell-derived EVs in treating ischemic stroke was cleared for clinical trials [7].

All EV therapies, regardless of their source, are subject to manufacturing challenges which influence their scalability and reproducibility [8, 9]. For example, EV yield, purity, and biological activity are dependent upon the isolation processes used [10]. Common methods, such as ultracentrifugation (UC) and ultrafiltration (UF), are subject to inefficiencies including variable EV yields, structural damage, aggregation, and non-EV protein contamination [11, 12], which may confound data and compromise the therapeutic efficacy of EV products [13]. Oscillation-based isolation (OSC) is a novel isolation technique which may enhance EV quality and quantity recovered, but is yet underexplored and must be evaluated against established methods [14]. As a key step in the EV manufacturing pipeline, advancing isolation processes will support EV production outcomes.

EV storage method affects downstream quality attributes and therapeutic viability [15]. Cryopreservation at -80°C is a conventional EV storage approach [16]; however, this method imposes logistical burdens for distribution of EV therapeutics due to its reliance on the pharmaceutical cold supply chain [17]. Lyophilization, or freeze-drying, has emerged as an attractive alternative for preserving EV integrity at ambient temperatures, eliminating cold storage needs and better supporting patient access as EVs progress to market [18]. Previous research has demonstrated that lyophilization successfully stabilizes EV preparations to varying levels of success dependent upon storage buffer/excipient application [19–21]. Additionally, lyophilized human platelet derived EVs are being applied in US FDA cleared clinical trials (<u>NCT04664738</u>), demonstrating the viability of this storage approach [22].

Evaluating the impact of EV isolation and storage conditions and elucidating the interplay between them will enhance manufacturing outcomes and accelerate the development of EV therapeutics [23]. This study investigates how EV isolation and storage techniques impact yield, structural integrity, purity, and potency. Our approach included nano-flow cytometric phenotyping to distinguish total nanoparticles from membrane-bound EV (Memglow^+^) and CD63^+^ EV portions. We calculated particle-to-protein ratios – a widely-used surrogate for EV purity [24–26] – to determine the relative purification performance of isolation methods. To compare preservation approaches, we assessed the NSC-EVs following storage via nano-flow cytometry, transmission electron microscopy, and a novel microglia morphology assay. This potency assay assesses a therapeutically-relevant function of NSC-EVs in the context of neurodegenerative diseases, as microglia play a key role in mediating neuroinflammation [27].

## MATERIALS AND METHODS

### Neuroepithelial cell culture and conditioned media collection

NSC-derived EVs were generated from WA09 human pluripotent stem cells (hPSCs) differentiated to neuroepithelium, as described in Lippmann et al. 2014 and illustrated by the timeline in Supplemental Figure 1. On day -1, hPSCs were seeded on Matrigel-coated 6-well plates at 200,000 cells/cm^2^ in E8 medium, and switched to E6 medium on day 0; both media were prepared in-house [28]. Fresh media changes were performed daily, and conditioned media were collected daily on days 4 through 9, shipped on dry ice, and stored at -80°C until processing. Cell identity was validated by immunostaining and confocal microscopy, with PAX6 and N-Cadherin (CHD2) stains being used to confirm neural progenitor identity (Supplemental Figure 1).

### Neural EV isolation and purification

Before purification, conditioned media were pooled into a batch and filtered through a 0.22-μm vacuum-filtration system (MilliporeSigma, Saint Louis, MO, USA), to remove cellular debris; filtered conditioned media (CM) also served as a control in comparing isolation methods. Each method was evaluated with at least six batches of NSC conditioned media.

#### OSC: oscillation-based isolation

The EXODUS H-600 Automatic Exosome Isolation System was used according to manufacturer instructions [14]. Conditioned media were processed using the high-purity setting in a large 30 nm pore-size EID cartridge (EXODUS Bio, CA, USA). EV concentrate was washed from the dual-filter nanopore membranes with 0.5 mL Bovine Serum Albumin Trehalose (BSAT) storage buffer (described below) and collected from the sample reservoir within the EID for storage and analysis.

#### UC: ultracentrifugation

Conditioned media were processed using a modified two-step ultracentrifugation protocol [29, 30]. The first round of UC was conducted at an RCFmax of 133,900 *× g* (Sorval WX UltraCentrifuge, ThermoFisher, MA, USA; Fiberlite F37L-8x100 Fixed-Angle Rotor, ThermoFisher, MA, USA) for either 1 hour (UC1) or 2 hours (UC2) at 4°C. The pellet was resuspended in cold PBS++, transferred to a micro-ultracentrifuge tube (CAT #54-361, ThermoFisher, MA, USA) and subjected to a second ultracentrifugation at 140,000 *× g* (Sorvall MX 150+ Micro-Ultracentrifuge, ThermoFisher, MA, USA; S110-AT Fixed-Angle Rotor, ThermoFisher, MA, USA) for 1 hour at 4°C. The final pellet was resuspended in BSAT storage buffer as described below to generate 0.5 mL aliquots for storage and analysis.

#### UF: ultrafiltration

Conditioned media was processed using a 100-kDa Amicon centrifugal filter unit (CAT #UFC905008, MilliporeSigma, MA, USA). The sample was washed with PBS++ and centrifuged at 4,000 *× g* for 10 minutes three times [31]. Following media filtration (3×), the filter unit was washed with PBS++ and the EV concentrate was collected in BSAT storage buffer as described below to generate 0.5 mL aliquots for storage and analysis.

### Neural EV storage

EV concentrate was diluted in a preservation buffer (described below); equal volumes of this EV isolate were subjected to cryopreservation and lyophilization for each isolation group. Buffer formulation and lyophilization protocols were adapted from Gorgens et al., 2022 and El Baradie et al., 2020.

#### EV preservation buffer formulation

The BSAT (BSA + trehalose) lyoprotectant buffer consisted of 0.2% bovine serum albumin (BSA; CAT #700-100P, Gemini Bio, CA, USA) and 25 mM trehalose (CAT #T5251, Sigma-Aldrich, MO, USA) in PBS++ [32], applied to EV concentrate at a 1:4 ratio.

#### Cryopreservation and thaw

EV isolates (0.5 mL) were aliquoted into a 1.5 mL microcentrifuge tube and rapidly frozen in liquid nitrogen for each isolation batch and stored at -80°C for one day. Cryopreserved samples were thawed on ice and vortexed briefly prior to analysis.

#### Lyophilization and rehydration

EV isolates (0.5 mL) were placed in a 1.5 mL microcentrifuge tube having a hole in the top and rapidly frozen in liquid nitrogen prior to lyophilization for each isolation preparation. Frozen isolates were lyophilized in a vacuum (-80°C, 0.035 mBar) overnight using a FreeZone Benchtop Freeze Dry system (Labconco, MO, USA). Tubes were recapped in aseptic conditions and stored at room temperature for one day. To rehydrate, the lyophilized samples were resuspended with distilled water to the original volume, vortexed briefly, and kept on ice until the point of analysis [32, 33].

### Analytical characterization

#### Size, count, and population analyses: nano-flow cytometry

EV isolates were diluted with PBS++ and evaluated on a NanoFCM Flow NanoAnalyzer (Xiamen, China) calibrated with 200-nm polystyrene beads and a silica nanosphere cocktail per manufacturer’s instructions. Particle concentration and size distribution were determined based on the flow rate and scattering intensity relative to the standard beads. To control for nanoparticle contributions from the formulation buffer, BSAT-only samples were included as blanks and used for background correction of EV isolate measurements. Additionally, population composition analysis was performed by incubating 20 uL of EV isolate with 1 µL of 100*×* diluted PC5-A human CD63 antibody to a final concentration of 2000*×* (CAT #BDB561983, BD Biosciences, NJ, USA) at 37°C for 2 hours in the dark. In the final 30 minutes, 1 µL of 100*×* diluted FITC Memglow (CAT #NC1950107, Cytoskeleton, CO, USA) was added. The double-positive subpopulation was identified as CD63+ EVs; nanoparticles possessing a lipid-membrane and CD63 tetraspanin protein. This nano-flow cytometry protocol was adapted from existing publications [34, 35], although these staining procedures were optimized based on a series of titration and incubation experiments.

#### Protein quantification: microBCA assay

Protein quantification was performed with EV concentrate immediately following isolation, prior to the addition of BSAT for storage, using the microBCA assay (Cat #23235, ThermoFisher, MA, USA) according to manufacturer’s instructions. EV concentrate was diluted 3 to 150*×* in duplicate to fall within the standard curve range (0 – 200 μg/mL BSA in PBS++). Diluted concentrate (75 µL) was mixed with reagent (75 µL), incubated for 2 hours at 37, and absorbance was read at 562nm using a spectrophotometer (Molecular Devices, CA, USA).

#### Single EV detection: TEM

OSC isolates were imaged using transmission electron microscopy (TEM), with a protocol adapted from Théry et al. 2006. A drop of isolate was deposited on carbon and Formvar-coated 400 mesh grids and incubated for 5 minutes to allow adsorption. The grids were rinsed with water and negatively stained with aqueous 0.5% w/v uranyl acetate. Excess solution was removed with filter paper, and images were acquired on a Gatan K3 IS direct electron detector (Gatan, Inc., CA, USA).

### In vitro microglia morphology assay

The microglia morphology assay was performed as previously described by Daga et al. (2024), using C20 (immortalized human microglia) cells in a 96-well plate format. Briefly, cells were stimulated using a cytokine cocktail of 5 ng/mL IFN-ү (Cat #PHC4033, Gibco, ThermoFisher, MA, USA) and 5 ng/mL TNF-α (Cat #10602HNAE, Sino Biological, Beijing, China). Treatment groups received the cytokine cocktail + 10 µL of cryopreserved or lyophilized OSC isolate at doses of 20,000, 200,000 or 2 million Memglow+ EVs/cell. Control groups received an equivalent dilution of BSAT buffer to match treatment conditions, thereby controlling for potential effects of BSA and trehalose on microglia morphology. After 24 hours of incubation, cells were fixed and imaged. Single cell morphological profiling was conducted via CellProfiler (Cambridge, MA, USA) using a pipeline developed by Daga et al. (2024). Twenty-one cellular and nuclear features were evaluated to assess global changes in microglia morphology, and this was plotted as a principal component analysis (PCA). This experiment was conducted with four batches of NSC conditioned media.

#### Radar Charts

Radar charts were generated in Microsoft Excel, to reflect both technical outcomes and practical considerations relevant to EV manufacturing. Yield and purification metrics were assigned scores based on groupings derived from statistical tests described below converted to ordinal scores using a predefined scale. When no statistically significant differences were observed between methods, all groups were assigned an equivalent score of 3. To preserve relative differences in performance between isolation methods, scores for concentrating effect were assigned using min-max normalization to a 1-5 range. Input volume compared the volume capacity of each method, with higher values indicating a larger starting volume of media processed within the same amount of time. Timeliness and ease scores are quantified such that higher values indicate faster and more user-friendly protocols.

### Statistical Analyses

The isolation and storage portion of this study was designed as a factorial experiment (4 × 2 design), in which EVs were isolated using four different methods (ISO) and stored under two conditions (STOR). The batch of NSC conditioned media served as the experimental unit, enabling evaluation of main effects of isolation and storage, as well as potential interactions between these factors. Purity metrics, expressed as particle-to-protein ratios, were log - transformed prior to analysis to stabilize variance and enable meaningful comparison across isolation methods. The in vitro experiment comparing storage conditions was conducted using a completely randomized design (CRD). Statistical analyses were performed using GraphPad Prism 10 (San Diego, CA, USA). Statistical significance was set at *P* < 0.05. One-way or two-way analysis of variance (ANOVA) with Tukey’s post hoc test was used for normally distributed data; non-parametric data were analyzed using the Kruskal-Wallis test followed by Dunn’s test with Bonferroni correction. Data are shown as mean ± SD or as box-and-whisker plots (min-max) with individual replicates.

## RESULTS

### Batch-to-batch variability differs between nanoparticle subpopulations

Conditioned media (CM) were collected in pooled batches from neuroepithelial differentiations of WA09 hPSCs [28]. Each cultures’ composition of PAX6+ neuroepithelial cells organized into polarized CDH2+ rosettes was qualitatively validated using immunocytochemical staining (Supplemental Fig. 1). Batch-to-batch variability in CM was assessed across five production batches, with Memglow^+^ EVs comprising 3.95% to 7.75% of total nanoparticles (Table 1). The coefficient of variation (CV) for Memglow^+^ EVs was 27.18%, indicating moderate variability across batches. CD63^+^/Memglow^+^ EVs represented a smaller proportion of total nanoparticles (0.32 to 1.43%), with greater batch variability (CV = 62.20%).

**Table 1.** Subpopulation analysis of five conditioned media (CM) production batches.

| Conditioned Media Batch |  |  |  |  |  |  |  |  |
| --- | --- | --- | --- | --- | --- | --- | --- | --- |
| Population | 1 | 2 | 3 | 4 | 5 | Standard deviation | Median absolute deviation | Coefficient of variation (%) |
| Memglow<sup>+</sup> EV<sup>†</sup><br>(% of total) | 7.57 | 7.75 | 5.24 | 3.95 | 5.45 | $1.63 \times 10^{-2}$ | $1.50 \times 10^{-2}$ | 27.18 |
| CD63<sup>+</sup>/Memglow<sup>+</sup> EV<sup>†</sup><br>(% of total) | 0.60 | 1.05 | 0.39 | 1.43 | 0.32 | $4.71 \times 10^{-3}$ | $2.80 \times 10^{-3}$ | 62.20 |

<sup>†</sup> Batch-to-batch variability in Memglow<sup>+</sup> extracellular vesicle (EV) content and Memglow<sup>+</sup>/CD63<sup>+</sup> EV content is shown as a percentage of total nanoparticles.

### Isolation method affects concentration measures but not size distribution or population composition of samples

Nano-flow cytometry analysis revealed similar nanoparticle size distributions across isolation methods (*P* > 0.05; Fig 1A); representative histograms can be viewed in Supplemental Fig. 2. Across isolation methods, nanoparticle size distributions exhibited a primary peak centered around 60 nm. There was no isolation method effect observed in either the proportion of Memglow+ EVs (P > 0.7487) or the proportion of CD63+/Memglow+ EVs (P > 0.4865) as a percent of total nanoparticles (Fig. 1B–C). Representative membrane labeling profiles measured by nano-flow cytometry can also be viewed in Supplemental Figure 2. Isolation method significantly affected nanoparticle concentration, Memglow^+^ EV concentration, and CD63^+^/Memglow^+^ EV concentration (*P* ≤ 0.0126; Fig. 1D–F). Isolation by UF yielded a greater (*P* = 0.0088) nanoparticle concentration than UC1, a greater (*P* < 0.0373) Memglow^+^ EV concentration than both UC1 and UC2, and a greater (*P* < 0.0003) CD63^+^/Memglow^+^ EV concentration than all other groups. No other isolation method differed significantly from another or from CM across any concentration metric (*P* > 0.1782; Fig. 1D–F).

**Figure 1.**
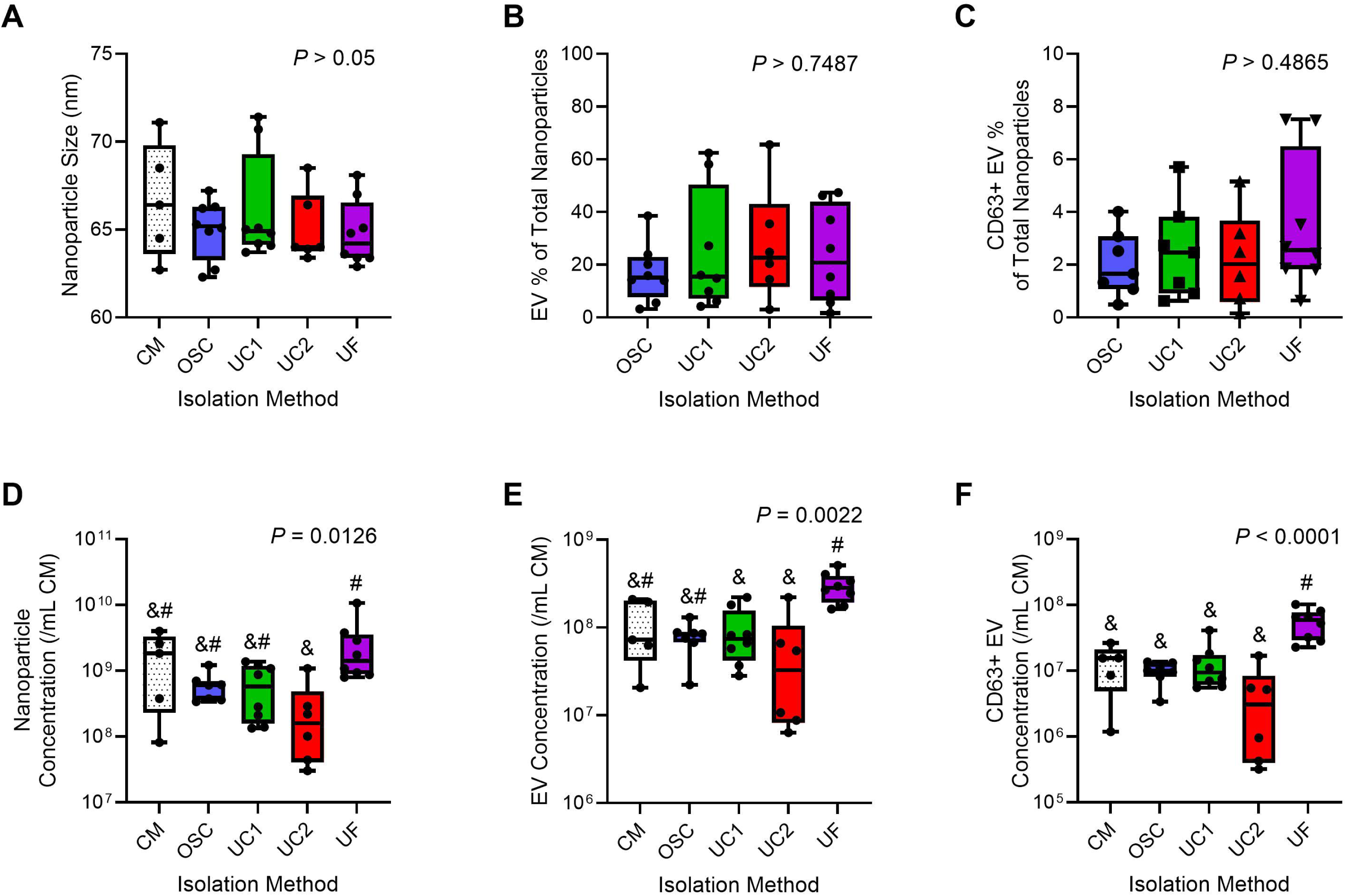
Nanoparticle size, population analysis, and concentrations across isolation methods. Panel A shows size distribution of nanoparticles, panels B and C show subpopulation composition, and panels D to F show concentration measurements for nanoparticles, EVs, and CD63^+^ EVs across isolation methods and conditioned media. **CM**, conditioned media; **OSC**, oscillation-based isolation; **UC1**, 1-hour ultracentrifugation; **UC2**, 2-hour ultracentrifugation; **UF**, ultrafiltration. (A) Mean nanoparticle size (nm) for each method. (B) Percent of the total nanoparticle population that comprises Memglow^+^ EVs, and (C) CD63^+^/Memglow^+^ EVs. (D–F) Concentrations of total nanoparticles, EVs (Memglow^+^), and CD63^+^ EVs (CD63^+^/Memglow^+^) are normalized to 1 mL of conditioned media to control for differences in input and output volumes across isolation methods. Within each panel, groups sharing a common symbol are not significantly different, whereas groups with different symbols differ significantly (*P* < 0.05), as determined by one-way ANOVA or Kruskal-Wallis tests with appropriate multiple comparisons corrections.

### Isolation method differentially affects protein content and purity of samples

There was an isolation method effect on total protein content (*P* < 0.0001) of samples and consequently on purity, calculated as a ratio of particles to µg of protein (Fig 2A–D). CM contained more (*P* < 0.0043) free protein than all isolated samples (Fig. 2A). Among isolation methods, samples isolated by UF contained more (*P* < 0.0001) free protein than all other groups, which did not significantly differ from one another (*P* > 0.3810; Fig. 2A). Isolation method significantly affected nanoparticle purity, Memglow^+^ EV purity, and CD63^+^/Memglow^+^ EV purity (*P* ≤ 0.0004) (Fig. 2B–D). OSC was the only method that produced a significant increase in purity relative to CM (*P* = 0.0102), whereas no other isolation method differed significantly from CM (*P* > 0.1762). Purity metrics did not differ significantly among the isolation methods (P > 0.2248; Fig. 2B–D).

**Figure 2.**
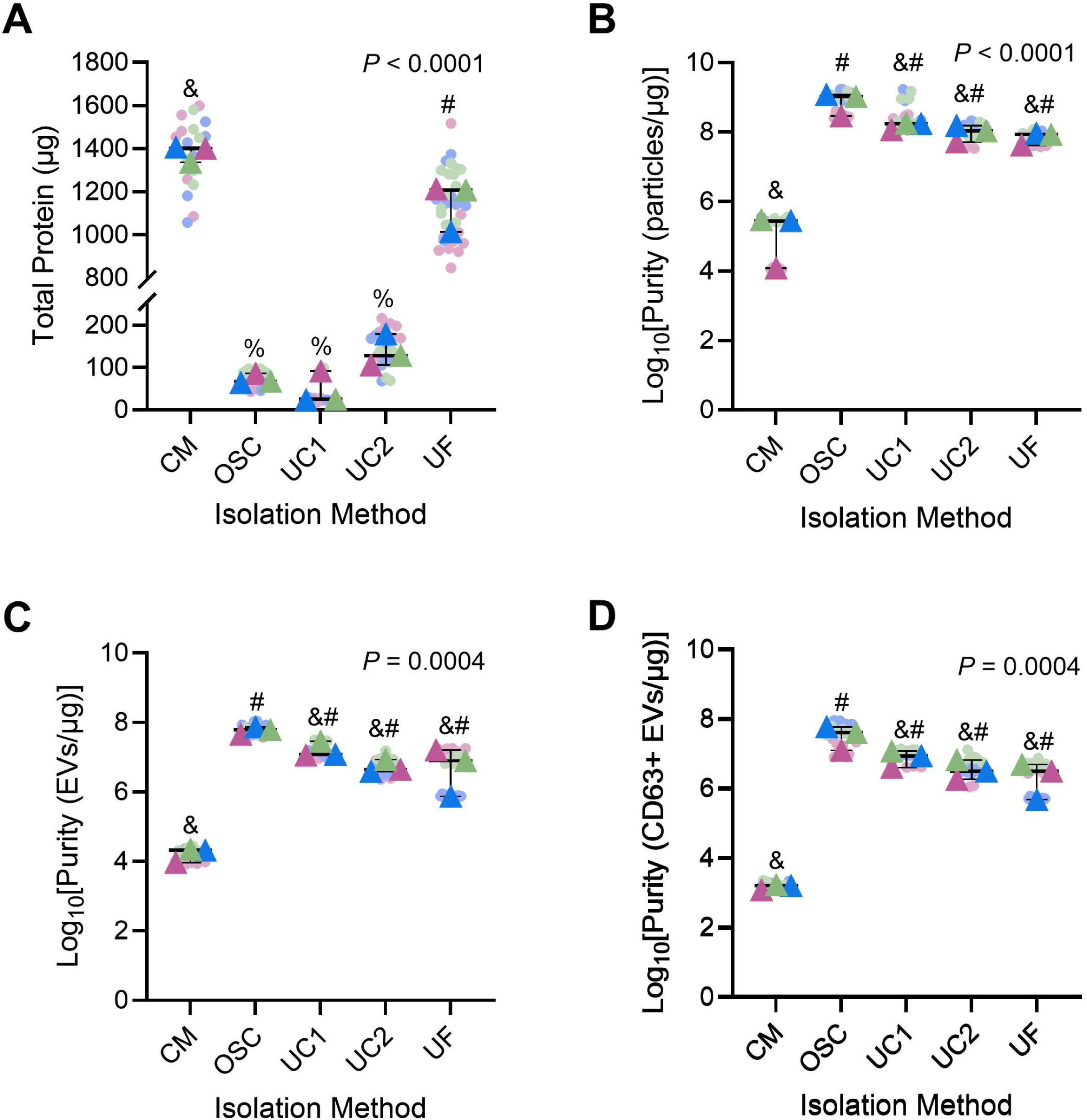
Effects of isolation method on protein content and purity of nanoparticles, EVs, and CD63^+^ EVs. (A) Total protein yield (µg) of isolated samples and conditioned media measured using a microBCA assay. (B–D) Purity of nanoparticles, EVs (Memglow^+^), and CD63^+^ EVs (CD63^+^/Memglow^+^), expressed as the ratio of particles to protein; purity values are displayed on a log□□ scale to facilitate comparison across orders of magnitude. Small circles represent technical replicates, and large triangles indicate experimental replicates (conditioned media batches). **CM**, conditioned media; **OSC**, oscillation-based isolation; **UC1**, 1-hour ultracentrifugation; **UC2**, 2-hour ultracentrifugation; **UF**, ultrafiltration. Within each panel, groups sharing a common symbol are not significantly different, whereas groups with different symbols differ significantly (P < 0.05), as determined by one-way ANOVA or Kruskal-Wallis tests with appropriate multiple comparisons corrections.

### Isolation methods exhibit distinct trade-offs across performance and workflow metrics

Radar charts were used to visualize performance of isolation methods based on yield, concentrating effect, and protein clearance for nanoparticles, Memglow^+^ EVs, and CD63^+^/Memglow^+^ EVs (Fig. 3A–C), as well as operational factors including input volume, timeliness, and ease (Fig. 3D). OSC exhibited strong performance in protein clearance and practical metrics, prompting its application for further evaluation of storage methods. UF performed well in yield and concentrating effects, and UC-based isolation approaches displayed intermediate profiles. This underscores trade-offs between yield, purification, and workflow considerations.

**Figure 3.**
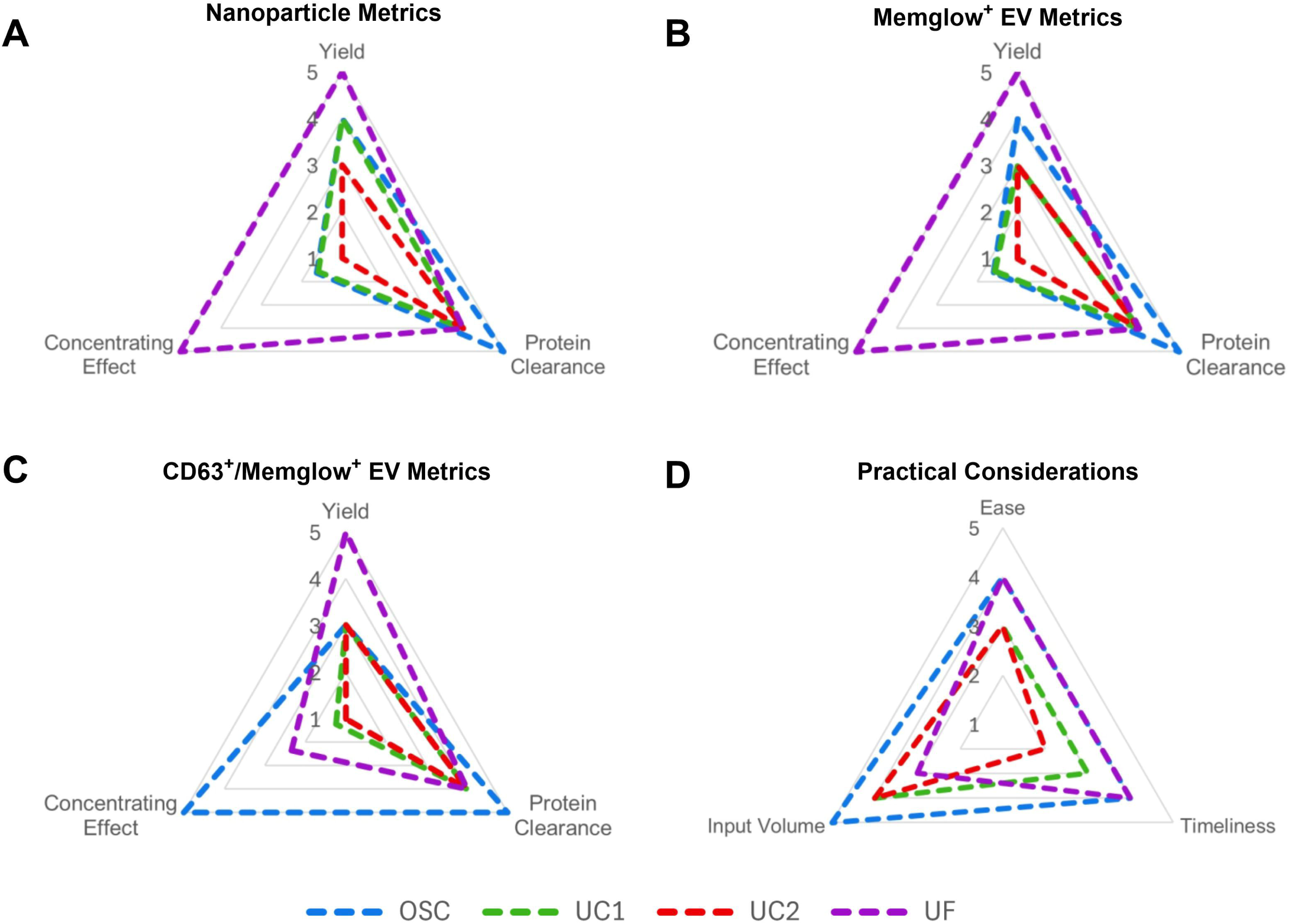
Relative efficiency of isolation methods in comparative radar charts. To facilitate integrated comparison of technical and practical performance, scaled radar charts summarize normalized, relative performance across metrics for each category of interest: (A) nanoparticles, (B) Memglow^+^ EVs, (C) CD63^+^/Memglow^+^ EVs, and (D) practical workflow considerations. For all charts, greater scores on the 1-5 scale indicate better relative performance within a given metric. Yield and protein clearance metrics were converted to ordinal scores based on statistically defined groupings, with a score of 3 assigned when no significant differences were observed among methods. Concentrating effect scores were normalized to a 1-5 range using min-max scaling to preserve relative differences between methods. Greater practical workflow scores correspond with larger starting volumes of conditioned media, faster processing times, and more user-friendly protocols.

### Lyophilization maintains size, population composition, and structure comparable to cryopreservation

Representative nanoparticle size distributions were comparable between storage conditions, with both cryopreserved (Fig. 4A) and lyophilized (Fig. 4D) nanoparticles exhibiting a unimodal peak centered around 60 nm. Nano-flow cytometry analysis revealed similar membrane labeling profiles between storage conditions, as demonstrated by the representative subpopulation scatterplots. The representative cryopreserved sample contained 38.4% Memglow^+^ EVs (Fig. 4B) and the lyophilized sample contained 35.6% Memglow^+^ EVs (Fig. 4E). Further characterization showed 2.1% of cryopreserved EVs were CD63^+^ (Fig. 4C) and 1.6% of lyophilized EVs were CD63^+^ (Fig. 4F). TEM imaging confirmed the presence of intact, spherical EVs under all storage conditions, with no major morphological differences observed between fresh (Fig. 4G), cryopreserved (Fig. 4H), and lyophilized (Fig. 4I) EVs, as depicted in the representative images.

**Figure 4.**
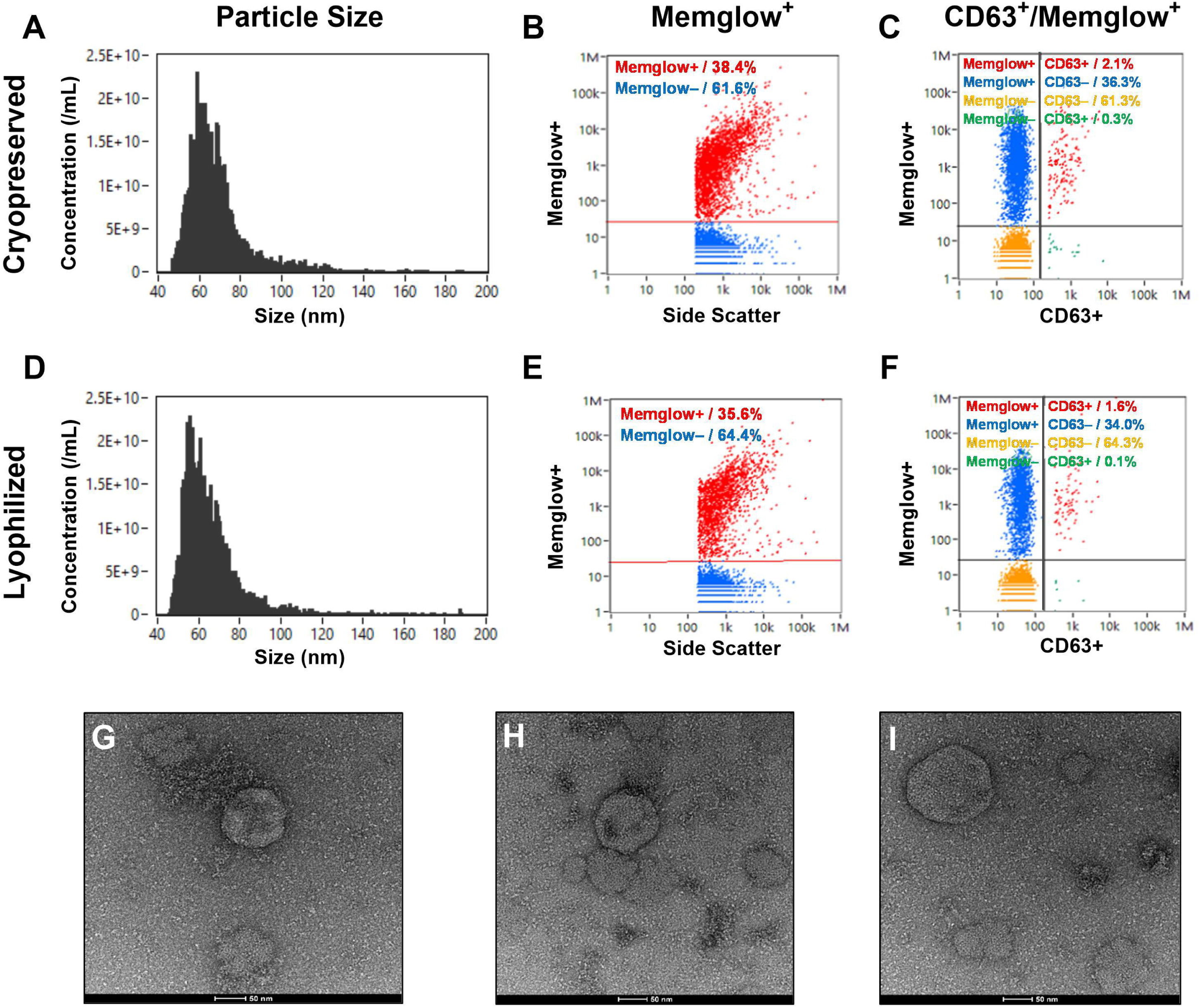
Nano-flow cytometry and transmission electron microscopy (TEM) analysis of OSC-isolated cryopreserved and lyophilized EVs. Panels A to C represent cryopreserved samples, while panels D to F represent lyophilized samples. (A, D) Size distributions of nanoparticles in each storage condition. (B, E) Scatter plots distinguishing Memglow^+^ (red) and Memglow^−^ (blue) populations. (C, F) Quadrant analysis of Memglow and CD63 labeling, identifying CD63^+^/Memglow^+^ (red), CD63^−^/Memglow^+^ (blue), CD63^+^/Memglow^−^ (green), and CD63^−^/Memglow^−^ (yellow) subpopulations. (G–I) Representative TEM images of OSC-isolated EVs under fresh (G), cryopreserved (H), and lyophilized (I) conditions. Scale bars = 50 nm.

### Size and concentration metrics do not differ between cryopreserved and lyophilized samples

No interaction between isolation method and storage condition was observed for any nano-flow cytometry metric (*P* ≥ 0.4488; Fig. 5A–D). A main effect of storage condition was detected for nanoparticle size (*P* = 0.0035); however, no significant differences were observed between cryopreserved and lyophilized sample pairs within any individual isolation method (*P* ≥ 0.3050; Fig. 5A). Storage condition did not significantly affect nanoparticle concentration, Memglow^+^ EV concentration, or CD63^+^/Memglow^+^ EV concentration (P ≥ 0.2618), and no differences were detected between cryopreserved and lyophilized samples within any isolation group (Fig. 5B–D).

**Figure 5.**
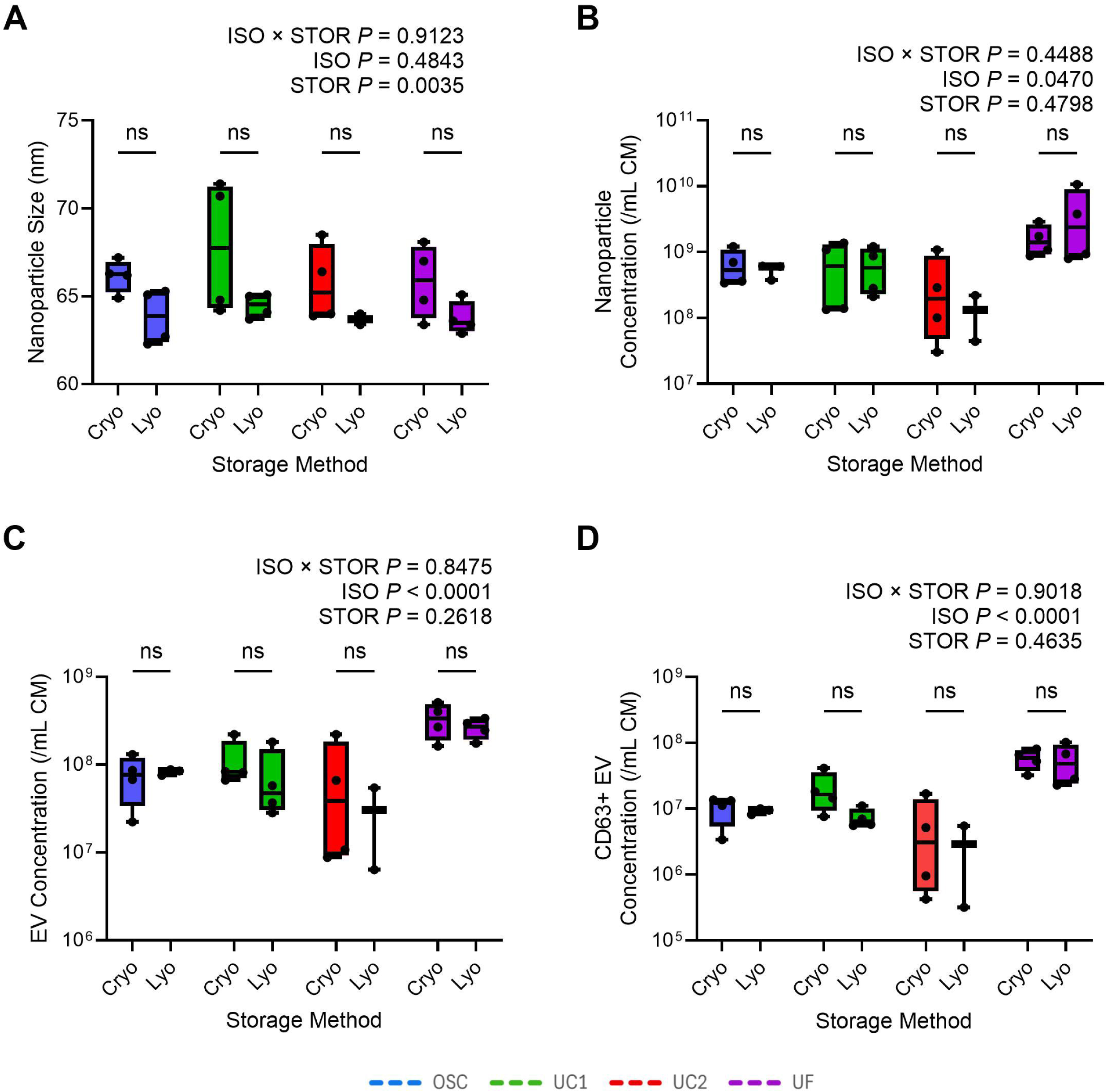
Nano-flow cytometry analysis of cryopreserved and lyophilized EVs within each isolation method. (A) Mean nanoparticle size (nm). (B–D) Concentrations of total nanoparticles, EVs (Memglow^+^), and CD63^+^ EVs (CD63^+^/ Memglow^+^), respectively, across isolation methods: **OSC**, oscillation-based isolation; **UC1**, 1-hour ultracentrifugation; **UC2**, 2-hour ultracentrifugation; **UF**, ultrafiltration. Two-way ANOVA with Tukey’s multiple comparisons was performed.

### Dose-dependent microglial immunomodulation by EVs is independent of storage method

Principal component analysis (PCA) was performed on 21 cellular and nuclear morphological features to evaluate microglial responses to OSC-isolated cryopreserved and lyophilized Memglow^+^ EVs (Fig. 6A). Representative images illustrate distinct morphological differences among unstimulated microglia (Fig. 6B), stimulated, untreated microglia (Fig. 6C), stimulated microglia treated with cryopreserved EVs (Fig. 6D), and stimulated microglia treated with lyophilized EVs (Fig. 6E). PCA component 1 (PC1) explained 74.8% of the variance (P < 0.0001) in this experiment, with no difference observed (*P* > 0.9999) between the immunomodulatory effects of cryopreserved and lyophilized EV treatments within each dose (Fig. 6F). A dose-dependent effect of EV treatment on PC1 was observed (*P* < 0.0001), with distinct microglia morphological changes compared to stimulated untreated microglia at 200,000 EVs/cell (*P* = 0.0002) and 2,000,000 EVs/cell (*P* < 0.0001; Fig. 6G).

**Figure 6.**
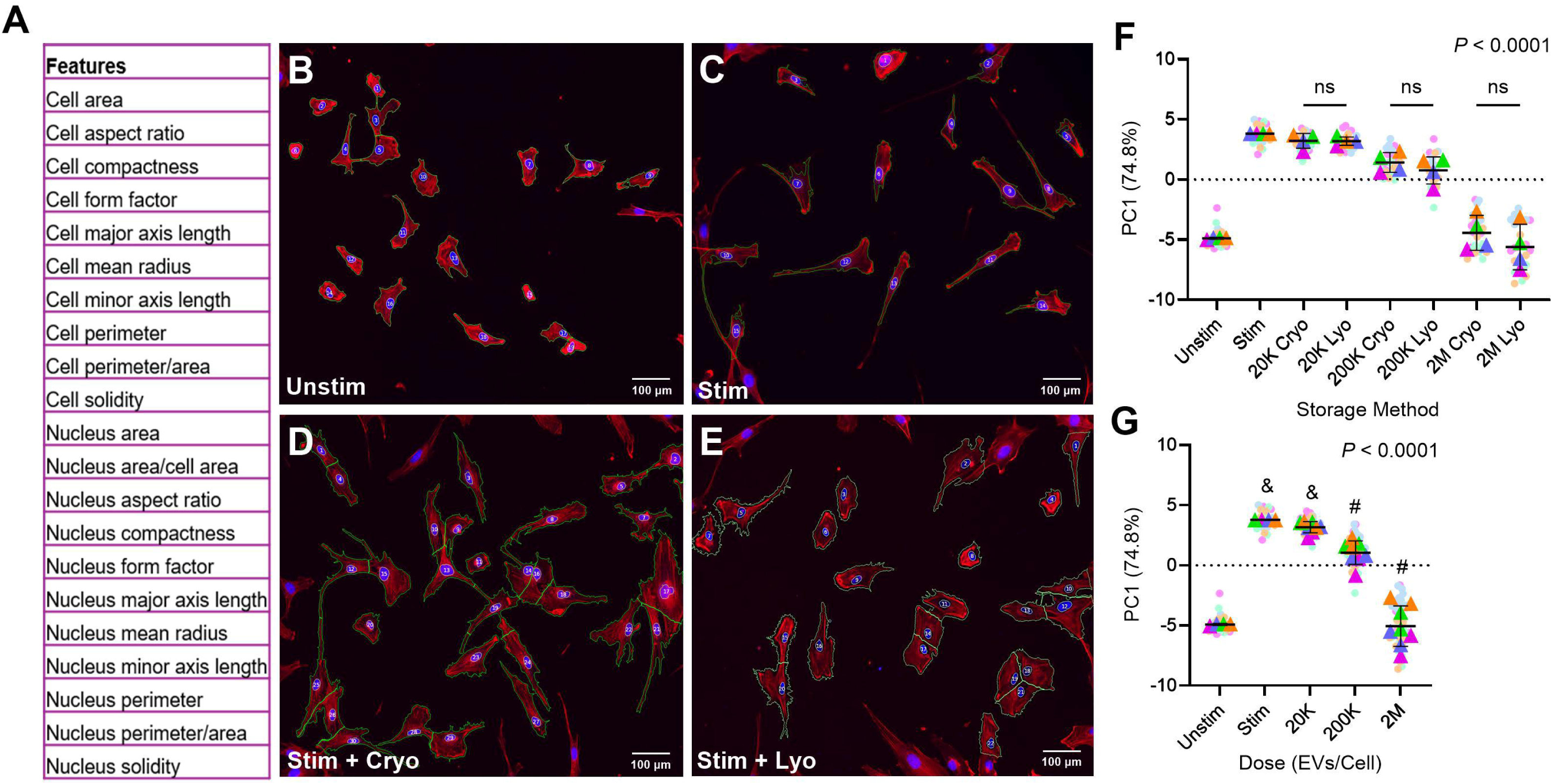
Immunomodulation of microglia morphology by OSC-isolated cryopreserved and lyophilized EVs. (A) List of 21 cellular and nuclear morphological features analyzed for principal component analysis (PCA). Representative images of (B) unstimulated microglia, (C) stimulated microglia without EV treatment, (D) stimulated microglia treated with cryopreserved EVs, and (E) stimulated microglia treated with lyophilized EVs. Scale bars = 100 µm. (F) Analysis of storage effect comparing impact of cryopreserved and lyophilized EVs within each dose level. (G) Analysis of dose-response comparing stimulated untreated microglia against EV-treated microglia to assess immunomodulatory effects; storage groups displayed in panel F are pooled to evaluate dose effect. PC1 accounts for 74.8% of the variance. Small circles represent technical replicates, and large triangles indicate experimental replicates (conditioned media preparations). **Unstim**, unstimulated; **Stim**, stimulated; **20K**, 20,000; **200K**, 200,000; **2M**, 2,000,000. Within each panel, groups sharing a common symbol are not significantly different, whereas groups with different symbols differ significantly (P < 0.05), as determined by one-way ANOVA or Kruskal-Wallis tests with appropriate multiple comparisons corrections.

## DISCUSSION

The manufacturing process is a critical determinant of extracellular vesicle (EV) quality and selecting appropriate processes may have implications for EV functionality [36, 37] This study evaluated the influence of both isolation method and storage approach on downstream product attributes. We compared four EV isolation methods and two storage conditions, focusing on yield, subpopulation composition, purity, and potency. We found key differences in method performance and implications for achieving scalable EV manufacturing workflows.

The efficiency and utility of EV isolation methods vary depending on factors including the sample source and EV subtype of interest [38]. Consequently, comparisons drawn from disparate studies – which often employ different isolation workflows, analytical pipelines, and biological systems – are inherently confounded, limiting standardization within the field [39, 40]. To address this challenge, we performed a controlled comparison of two widely adopted isolation approaches, ultracentrifugation (UC) and ultrafiltration (UF), alongside a less commonly utilized oscillation-based isolation method (OSC) using a single EV source and consistent analytical techniques. To simplify cross-method evaluation, we employed radar chart visualizations to integrate technical and practical performance metrics within a unified multi-dimensional framework [11]. Among the isolation methods tested in this comparison, (1) OSC was the only approach that achieved significant purification relative to conditioned media (Fig. 2); (2) UC, despite its widespread use, did not demonstrate purification beyond that of conditioned media, and required lengthy spin times and large starting volumes (Fig. 3); and (3) UF, while capable of modest EV enrichment, similarly did not achieve purification relative to conditioned media (Fig2).

Protein co-purification is a well-recognized limitation of EV isolation methods. Contaminating proteins and soluble factors can obscure EV-specific bioactivity and confound downstream analyses [39, 41–43]. Isolation method significantly influenced protein clearance, and consequently particle-to-protein ratios, which represent purification. The purification performance of oscillator-based isolation is consistent with prior reports suggesting that acoustic nanofilter approaches which rely upon oscillatory pressure effectively separate EVs from protein aggregates, cell debris, and other non-EV particulates [44–46]. These results also align with literature acknowledging UC-isolation limitations, in particular, non-EV protein contamination and potential structural damage from high g-forces [11].

Nano-flow cytometry was used to enable subpopulation-level characterization of the conditioned media particles isolated, including non-EV contamination. Unlike nanoparticle tracking analysis (NTA), which reports total nanoparticle counts without distinguishing EV identity, nano-flow cytometry enables phenotyping through detection of membrane and lumen markers [35]. EV isolates contain membrane-bound vesicles, among other biological nanoparticles, including lipoproteins, exomeres, supermeres, excipients, and protein aggregates; these non-vesicular constituents contribute to total particle counts but may differ substantially in composition and bioactivity [41, 47–49]. By identifying membrane-bound (Memglow) and CD63 tetraspanin-expressing subpopulations, nano-flow cytometry distinguished EVs from non-vesicular particles, revealing that EVs comprised only a fraction of total nanoparticles in conditioned media. In concordance with prior reports demonstrating morphological and molecular heterogeneity within EV isolates derived from a single cell type [50–53], we observed batch-to-batch variability in EV subpopulation composition (Table 1), underscoring the importance of interrogating EV identity with specificity rather than relying solely on bulk particle measurements. Phenotypic EV characterization therefore provides a basis for linking EV manufacturing parameters to product composition, offering a valuable tool for evaluating therapeutic EV yield and consistency.

An additional objective of this work was to compare conventional cryopreservation at - 80°C with lyophilization based on structural, phenotypic, and functional criteria. Lyophilization in a cryoprotectant buffer preserved the morphology, size distribution, and subpopulation composition of EVs at levels comparable to cryopreserved controls. Prior studies with bovine milk and mesenchymal stromal cell EVs support the use of BSA and trehalose (BSAT) as effective lyoprotectants [23, 32], and this study demonstrates the compatibility of these excipients with neural stem cell-derived EVs. Although we did not explore alternative excipient formulations, other groups have reported success with a variety of stabilizing agents and protocols for lyophilized EVs [20, 21]. Future work may expand upon excipient optimization for neural EV systems, with the goal of further enhancing the stability and product shelf life of this nanotherapeutic.

Lyophilization preserves therapeutic EVs and is used for EV human clinical trials [18, 54, 55]. The present study applied this approach for neural stem cell-derived EV stabilization. Lyophilized mesenchymal stromal cell sourced EVs [56, 57] and tumor-derived EVs [19, 58] retain their character and function both in vitro and in vivo, but these systems differ in their cellular origin and intended application. Here, we showed that lyophilized NSC-derived EVs retained their structural integrity and subpopulation composition, confirming that this preservation strategy is applicable beyond previously tested EV types.

Microglial morphology serves as an integrative proxy for cellular function and behavior, reflecting coordinated changes in inflammatory secretion, proteomic composition, and mitochondrial organization [27]. Accordingly, microglial morphology provides a meaningful assessment of EV-mediated immunomodulation. NSC-EVs modulate in vivo microglial activation and viability [6], with implications for treatment of neurodegenerative diseases. These EVs influence immune dampening and attenuate pro-inflammatory responses, promoting polarization toward an M2-like phenotype, thereby enhancing tissue repair and neuroprotection [59–61]. Consequently, NSC-EVs are of particular interest for treating acute brain injuries and neurodegenerative diseases [1–3, 62].

Lyophilized NSC-EVs demonstrated dose-dependent immunomodulatory activity in vitro, comparable to their cryopreserved counterparts (Fig. 6), supporting the deployment of lyophilized EV therapeutics in resource-limited or decentralized settings [18]. By pairing structural analyses with this phenotypic potency assay, the present study aligns with the field’s growing emphasis on robust, scalable in vitro assays as a foundation for therapeutic EV quality control, potency assessment, and product characterization [63–65]. Considering the natural heterogeneity of conditioned media and the influence of manufacturing processes on functional outcomes, in vitro potency assays are critical for ensuring quality control and consistency between batches and products [66, 67]. Additionally, by tailoring potency assays to the specific disease and proposed mechanism of action, as we have done in this study, it is possible to predict the therapeutic efficacy of an EV product and evaluate its effectiveness [63, 68].

This study demonstrates that the combination of various isolation and storage strategies influences manufacturing outcomes, with consideration for EV purity and functional integrity. Among the methods tested, oscillator-based isolation paired with lyophilization offers a promising, cold chain–independent manufacturing pipeline to produce therapeutic EVs. The ability to preserve EV integrity and potency under ambient conditions enables wider distribution of NSC-EV therapies [18]. These findings establish a framework for evaluating EV isolation methods within a controlled manufacturing context, and inform the development of scalable EV production pipelines tailored to specific cell sources and therapeutic applications.

## Author Contributions and Acknowledgements

Conceptualization, MEG and SLS; methodology and validation, all authors; investigation, MEG, LCM, KRD; data curation and formal analysis, MEG, LCM, KRD, FS; writing—original draft preparation, MEG and SLS; writing—review and editing, all authors; visualization, MEG, KRD, FS; supervision, project administration, and funding acquisition, SLS. All authors have read and agreed to the published version of the manuscript. The authors would like to acknowledge and express thanks for the support of the Georgia Research Alliance and the National Science Foundation Engineering Research Center for Cell Manufacturing Technologies (NSF Grant No. EEC-1648035), as well as Shanmathi Ramasubramanian for her help with statistical analysis.

---

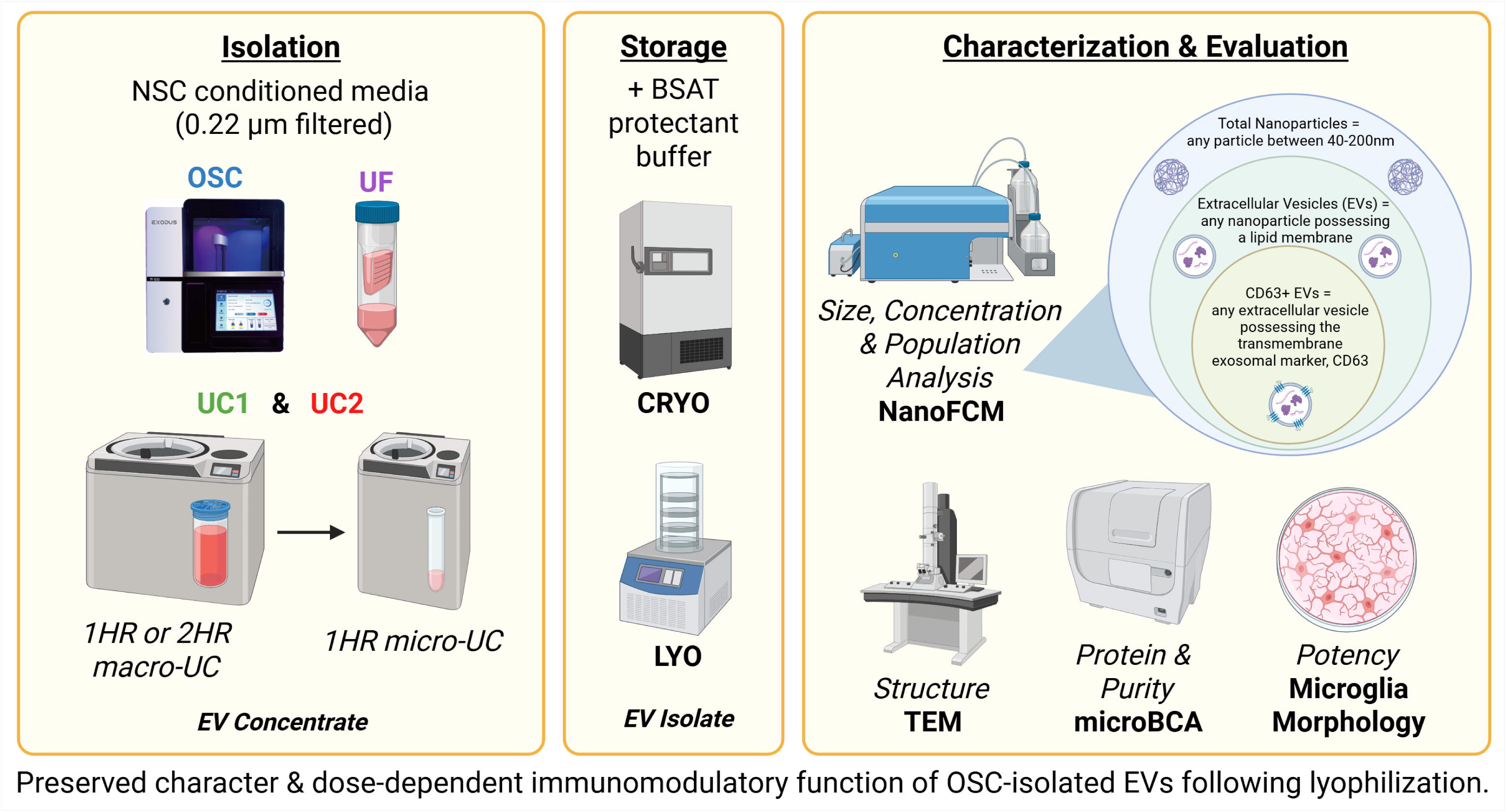

**Supplementary Figure 1. Cell culture timeline and confocal microscopy of immunostained cell culture platform verifying neural cell identity.** Human neuroepithelial cultures were immunostained to confirm neural progenitor identity prior to EV collection. (A) Red staining indicates PAX6, a marker of neural progenitor nuclei, while green staining shows N-cadherin (B), highlighting the polarized rosette structures characteristic of neuroepithelial organization; the merged image captures the overlap of these stains (C). Scale bars = 100 µm.

**Supplementary Figure 2. Nano-flow cytometry characterization of nanoparticle size distribution and sample subpopulations across four isolation methods.** NanoFCM Flow NanoAnalyzer outputs of groups, organized by row: BSA + trehalose buffer (**BSAT)**, conditioned media (**CM**), oscillation-based isolation (**OSC**), 1-hour ultracentrifugation (**UC**O), 2-hour ultracentrifugation (**UC**O), and ultrafiltration (**UF**) from top to bottom. The left column displays size distributions of nanoparticles in each group. The middle column shows side scatter plots distinguishing Memglow^+^ (red) and Memglow^−^ (blue) populations. The right column depicts quadrant analysis for Memglow and CD63 markers, differentiating Memglow^+^/CD63^+^ (red), Memglow^+^/CD63^−^ (blue), Memglow^−^/CD63^+^ (green), and Memglow^−^/CD63^−^ (yellow) subpopulations. Preserved character & dose-dependent immunomodulatory function of OSC-isolated EVs following lyophilization.

