## Supplementary figures and images for "Impact of Isolation and Storage Methods on the Properties of Neural Extracellular Vesicles"

### Supplemental Figure 1

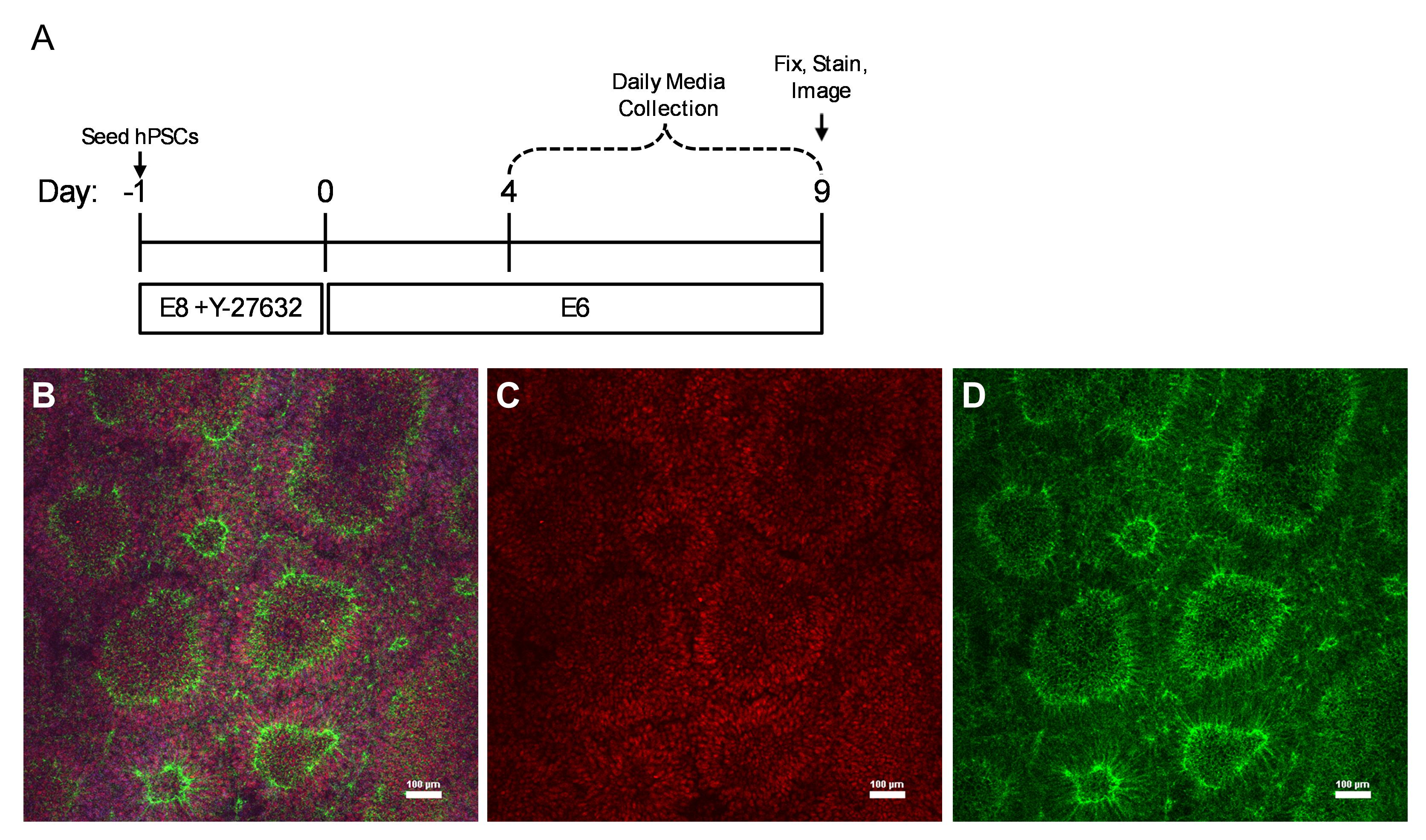

### Supplemental Figure 2

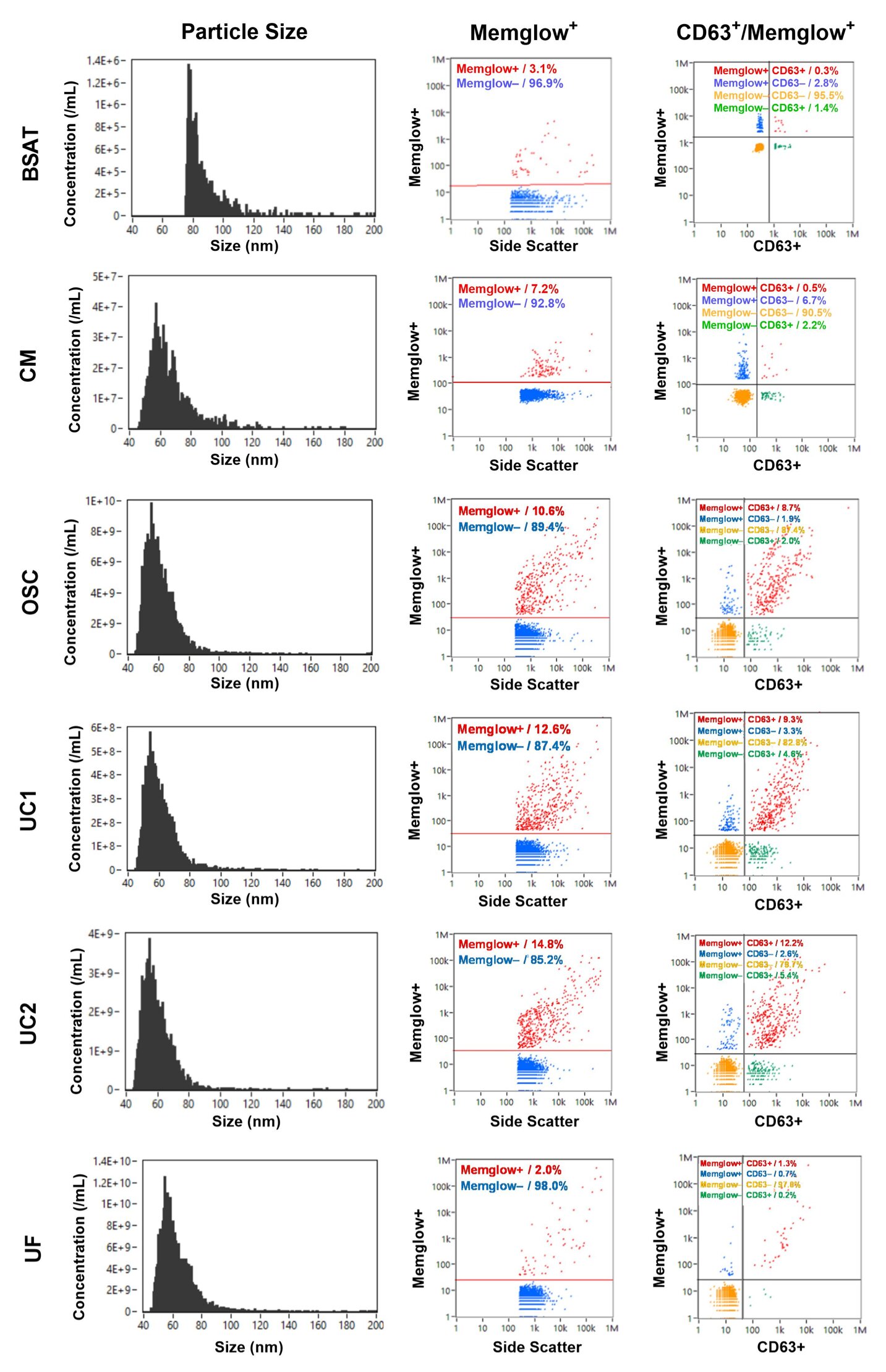
